# *Candida khanbhai* sp. nov., a new clinically relevant yeast within the *Candida haemulonii* species complex

**DOI:** 10.1101/2022.12.01.518802

**Authors:** Auke W. de Jong, Khaled Al-Obaid, Ratna Mohd Tap, Bert Gerrits van den Ende, Marizeth Groenewald, Leena Joseph, Suhail Ahmad, Ferry Hagen

## Abstract

Invasive fungal infections caused by non-*albicans Candida* species are increasingly reported. Recent advances in diagnostic and molecular tools enabled better identification and detection of these emerging pathogenic yeasts. Several of these emerging species belong to the *Candida haemulonii* species complex, which attracted much attention due to the rapid global emergence of its multi-drug resistant member *Candida auris*. Here, we describe a new clinically relevant yeast isolated from geographically distinct regions, representing the proposed novel species *Candida khanbhai*, member of the *C. haemulonii* species complex. Moreover, several members of the *C. haemulonii* species complex were observed to be invalidly described, including *Candida auris* and *Candida vulturna*. Hence, the opportunity was taken to correct this here, formally validating the names of various yeast species.

## Introduction

The *Candida* genus (*Ascomycota, Saccharomycotina, Saccharomycetes*, *Saccharomycetales*) contains some of the most well-known human fungal pathogens. Although *Candida albicans* remains the primary cause of candidiasis, infections caused by *non-albicans Candida* species are increasingly reported (1). Globalization, climate change, and a steadily growing population of immunocompromised patients over the last few decades paved the way for rare and new species to emerge as fungal pathogens (2–4). Recent advances in diagnostic and molecular tools enabled better detection and identification of these emerging fungi. Therefore, the list of reported species causing human infection is rapidly expanding.

Several of these emerging pathogens group within the *Candida haemulonii* species complex. Members of this complex belong to the family of the *Metschnikowiaceae* and are closely related to the genus *Clavispora* (5). Lately, the *C. haemulonii* species complex has drawn much attention because of its multi-drug resistant nature and the rapid worldwide spread of one of its members, *Candida auris* (6). Next to *C. auris* other members also cause nosocomial infections, including *Candida haemulonii*, *Candida duobushaemulonii*, *Candida pseudohaemulonii* and *Candida vulturna* (5,7). Some of these species have been isolated from ecological sources such as fish (*C. haemulonii*), flowers (*C. vulturna*) or insects (*C. duobushaemulonii*) (5,8). However, most reported strains have a human origin. For example, *C. pseudohaemulonii* and *C. auris* were described from clinical cases since no environmental strains were available (9,10). Only very recent sampling efforts linked *C. auris* to the marine ecosystem, but the ecological niche of *C. pseudohaemulonii* remains to be explored (11,12). In the present study, we describe a new yeast isolated from geographically distinct clinical samples, representing a proposed novel species, *Candida khanbhai,* related to members of the *C. haemulonii* species complex. Additionally, it was observed that several members of the *Candida haemulonii* species complex were invalidly described. Hence, the opportunity was taken to correct this here, formally validating the names.

## Case descriptions

Two clinical strains were obtained. Case 1: The strain (Kw2195/19 = CBS 16213^T^) was collected in 2019 from a 73-year-old male patient during a suspected *C. auris* outbreak involving two patients in the Mubarak Al-Kabir Hospital, Jabriya, Kuwait. Besides reinforcing infection control measures and isolation precautions, all patients in the same medical ward were screened for *C. auris* nasal carriage. The patient’s nasal swab was subcultured onto Sabouraud dextrose agar, which grew cream-colored colonies. Identification by VITEK MS (bioMérieux, Marcy-l’Étoile, France) could not reliably identify the strain. Instead, VITEK 2 (bioMérieux) was used. The result was indicative of *C. duobushaemulonii* (99% score). Of note, the patient did not receive antibacterial or antifungal agents in the last three months and was not infected or colonized with *Candida* species throughout his stay in the hospital. Given the discrepant identification results the strain was sent to the Westerdijk Fungal Biodiversity Institute for further analyses.

Case 2. The strain (UZ687/14 = CBS 16555) originated from the blood culture of a 55-year-old male patient admitted in July 2014 to the Tengku Ampuan Afzan Hospital, Pahang, Malaysia, due to having generalized body weakness, lethargy, nausea, and a slight fever of 37.8°C. Blood cultures taken on day one and day four were negative. On day nine, the patient developed a high-grade fever of 39.5°C. Chest X-ray showed pulmonary infiltration in the bilateral lower zone. The patient was treated for new onset hospital-acquired pneumonia. Intravenous tazocin (562 mg/6 hour) was given for seven days. The blood culture taken on day nine grew yeast-like cells that were identified biochemically by API 20C (bioMérieux) as *Clavispora lusitaniae* (81% score) after 72h of incubation. A dosage of 400 mg/day intravenous fluconazole was started at day 12. Nevertheless, the patient’s clinical condition deteriorated further, and he succumbed to the infection two days later. The strain was sent to the Westerdijk Fungal Biodiversity Institute for further analyses.

## Material & Methods

### DNA sequencing and phylogenetic analysis

After receiving both strains at the mycological reference center (Westerdijk Fungal Biodiversity Institute, Utrecht, the Netherlands) they were subcultured onto 2% Glucose, 0.5% Yeast extract, 1% Peptone, and 1.5% agar (GYPA) plates. After 48h of incubation at 35°C the strains were further processed for molecular characterization and culture collection deposition. DNA was extracted using a manual standardized cetyltrimethylammonium bromide-based method, as previously described (13). Similarly, this was done for a reference set of type-strains, representing all members of the *C. haemulonii* species complex: *C. auris* (CBS 10913^T^, clade II; AR0383, clade III; AR0385, clade IV; AR0387, clade I; AR1097, clade V), *Candida chanthaburiensis* (CBS 10926^T^), *C. duobushaemulonii* (CBS 7798^T^), *Candida heveicola* (CBS 10701^T^), *C. haemulonii* (CBS 5149^T^), *Candida konsanensis* (CBS 12666^T^), *C. pseudohaemulonii* (CBS 10004^T^), *Candida metrosideri* (CBS 16091^T^), *Candida ohialehuae* (CBS 16092^T^), *Candida ruelliae* (CBS 10815^T^), *C. vulturna* (CBS 14366^T^), and *C. lusitaniae* (CBS 6936^T^). Type strains were obtained from the CBS culture collection of the Westerdijk Fungal Biodiversity Institute, while most of the *C. auris* reference strains came from the CDC Isolate Bank (US Centers for Disease Control & Prevention, Atlanta, GA, USA).

Amplified Fragment Length Polymorphism (AFLP) fingerprint analysis was performed as previously described, except that the fragment analysis was performed using the ABI3700×L Genetic Analyzer platform (Applied Biosystems, Palo Alto, CA, USA) (14). Raw fingerprint data were analyzed in Bionumerics v7.6 (Applied Math, St. Martens-Latem, Belgium) and a dendrogram was created using the Pearson correlation similarity coefficient and UPGMA cluster analysis algorithms.

Amplification of the D1/D2 region of the large subunit (LSU) ribosomal RNA gene and the internal transcribed spacer ITS1-5.8S-ITS2 regions were done using the NL1 (5′-GCATATCAATAAGCGGAGGAAAAG-3′) plus NL4 (5′-GGTCCGTGTTTCAAGACGG-3′) and ITS1 (5′ TCCGTAGGTGAACCTGCGG 3’) plus ITS4 (5′ TCCTCCGCTTATTGATATGC 3′) primer pairs, respectively (15). Sequencing was performed with the same primers using BigDye chemistry v3.1 and sequences were generated onto the ABI3700xL Genetic Analyzer platform (Applied Biosystems). Raw data bi-directional sequences were manually checked and corrected using the SeqMan module of the Lasergene v17 software package (DNASTAR, Madison, WI, USA).

The aligned ITS (455bp) and LSU (535bp) datasets were concatenated in SequenceMatrix v1.8 (16). The concatenated sequences were aligned using MAFFT v7.409 with FFT-NS-i option (17). The best model of substitution was established by ModelFinder resulting in TNe+G4 according to Bayesian Information Criterion (BIC) (18). Phylogenetic trees were estimated using maximum likelihood with IQ-tree v1.6.12 (19) calculating likelihood ratio test, Bayes inference, and ultrafast bootstrap values running 1,000 bootstraps showing the values in this order on the nodes of the tree.

### Phenotypic/physiological characterization

The colony morphology of both strains was determined using plate cultures grown on GYPA for seven days at 25°C or up to one month at room temperature. Microscopic pictures were taken of cells grown in YPD broth (1% Yeast extract, 2% Bacto Peptone, 2% Glucose) at 200 rpm, 25°C for 24 hours, using an Axioskop 2 plus (Carl Zeiss, Jena, Germany) microscope fitted with a Nikon DS-Ri2 microscope camera (Nikon Instruments, Melville, NY, USA). Additionally, the mating compatibility of both strains was tested by mixing them onto GYPA and Malt-Extract Agar (MEA; Oxoid, Basingstoke, U.K.) plates and incubated at 25°C for up to 12 weeks. Given the close phylogenetic relatedness of the studied strains to *C. auris* and *C. haemulonii* (see results), the colony phenotype was compared to the type strains of *C. haemulonii* (CBS 5149^T^) and *C. auris* (CBS 10913^T^) on the new CHROMagar *Candida* Plus (CCP) (CHROMagar, Paris, France), specifically designed to differentiate *C. auris* from other pathogenic yeasts. Finally, (mis)identification of the strains by matrix-assisted laser desorption ionization–time of flight mass spectrometry (MALDI-TOF MS) was tested using a Bruker MALDI Biotyper system (database v11). MALDI-TOF MS main spectra (MSP) of both strains were generated using the manufacturers’ instructions. Physiological characterization of both strains was done using standard procedures previously described (20). Fermentation and assimilation of carbohydrates was performed in liquid media at 25°C up to 21 days. Assimilation of nitrogen compounds was assessed with the auxanographic method.

### Antifungal susceptibility testing

Due to the clinical origin of both strains and their relatedness to the multidrug resistant members of the *C. haemulonii* complex, their in vitro antifungal susceptibility was determined using the EUCAST protocol (21). Antifungals were obtained from Sigma-Aldrich (St. Louis, MI, USA). Included in this test were: Amphotericin B (AMB), anidulafungin (AND), micafungin (MCF), 5-flucytosine (5FC), and various azoles including fluconazole (FLU), itraconazole (ITR), voriconazole (VOR), posaconazole (POS) and isavuconazole (ISA). The concentrations tested ranged between 0.03 and 16 μg/mL for all drugs, except fluconazole (0.125–64 μg/mL). MIC values were determined after 24h and 48h.

## Results

### Identification and phylogeny

Molecular based identification of CBS 16213^T^ and CBS 16555 by ITS sequencing resulted in low-score NCBI Nucleotide database BLAST hits with a closest hit of 86% similarity to *C. duobushaemulonii*. Similarly, the D1/D2 sequence of these strains had both a similarity of 92% with *C. heveicola* using the NCBI Nucleotide database (Figure 1). By AFLP analysis CBS 16213^T^ and CBS 16555 shared an 84% fingerprint similarity, but had only 35% similarity compared to the fingerprints of *C. heveicola* and *C. chanthaburiensis* type strains and 24% similarity with *C. konsanensis* (Figure 1). Phylogenetic analysis of the concatenated ITS and D1/D2 sequence data showed that CBS 16213^T^ and CBS 16555 clustered tightly together and form a lineage that is basal to a group of eight *C. haemulonii* species complex members, of which three (*C. duobushaemulonii*, *C. pseudohaemulonii* and *C. vulturna*) are clinically relevant (Figure 2).

**Figure 1.**
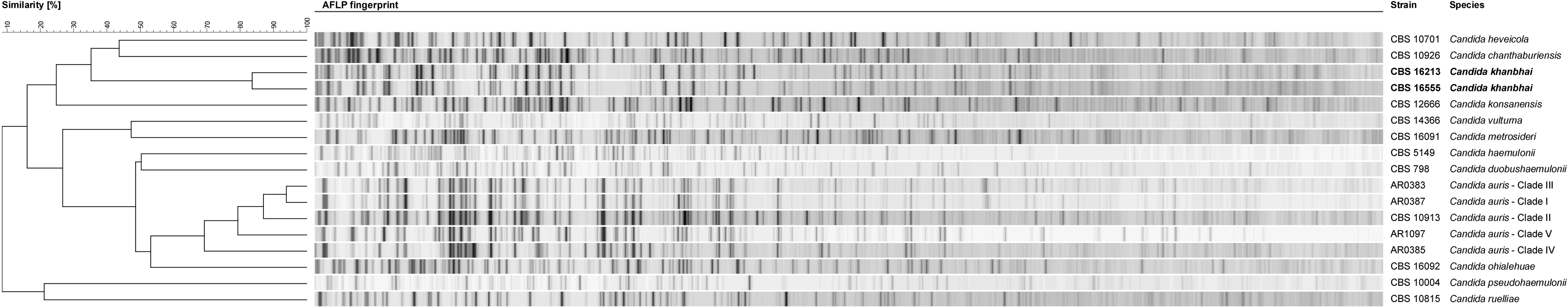
AFLP fingerprint analysis of the *Candida haemulonii* species complex. An AFLP fingerprint analysis was initially performed to investigate if both *Candida khanbhai* strains CBS 16213 and CBS 16555 (in bold) had similar fragment patterns as any of the other described species within the *Candida haemulonii* species complex. *C. khanbhai* strains CBS 16213 and CBS 16555 had an 84% similarity, but had only 35% similarity compared to the AFLP fingerprints of *Candida heveicola* and *Candida chanthaburiensis* type strains and 24% similarity with *Candida konsanensis.* Less than 20% similarity was observed with any of the other *C. haemulonii* species complex members.

**Figure 2.**
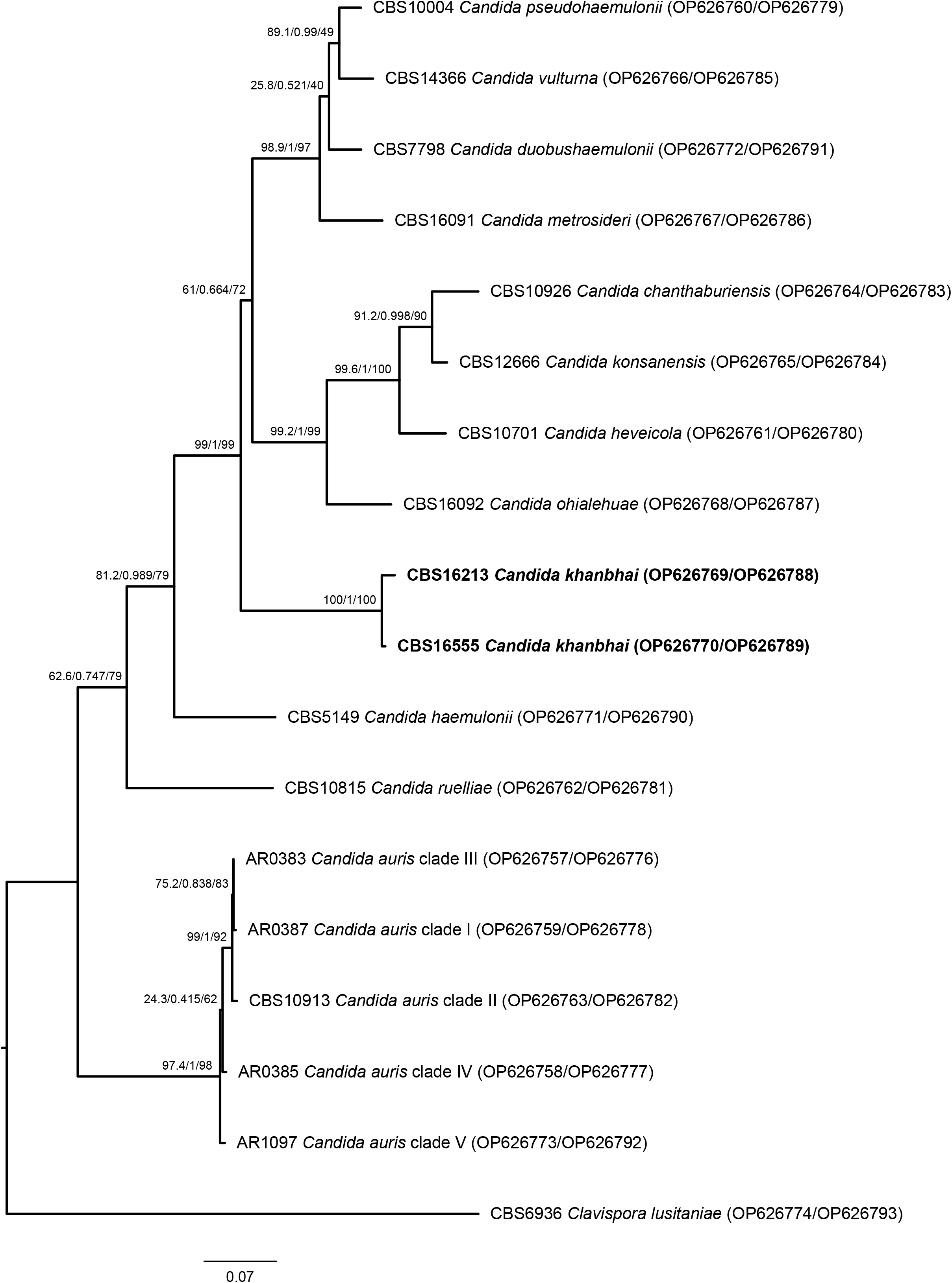
Phylogenetic analysis of the *Candida haemulonii* species complex. Sequenced based analysis was done by using the aligned and concatenated internal transcribed spacer (ITS1-5.8S-ITS2) region and the D1/D2 region of the large subunit (LSU) of the ribosomal DNA for the type strains representing the members of the *Candida haemulonii* species complex plus the here described *Candida khanbhai* strains CBS 16213^T^ and CBS 16555 (in bold). *Clavispora lusitaniae* (CBS 6936) was used as outgroup. The best model of substitution was observed to be TNe+G4 according to Bayesian Information Criterion (BIC) (18). Phylogenetic trees were estimated using maximum likelihood with IQ-tree v1.6.12 (19). The likelihood ratio test, Bayes inference, and ultrafast bootstrap values (1,000 bootstraps) are in this order placed on the nodes of the tree.

### Phenotypic characterization

Both clinical strains CBS 16123^T^ and CBS 16555 were morphologically very similar. Pictures of the type strain (CBS 16123^T^) are given in Figure 3 A-C. On GYPA plates, colonies are off-white coloured, glossy, soft, butyrous and with an entire margin and do not change morphology from 7 days up to 1 month of incubation (Figure 3A). After 24h in YPD broth, cells were ovoid, ellipsoidal to spherical (1.5–3.5 × 2.5–4.5 μm in diameter) (Fig. 3B). After seven days of growth on GYPA plates, cells are shaped similarly as in liquid culture, but the formation of a sub-population of enlarged cells that are up to 6 μm in diameter can be observed (Figure 3C). Cells occur singly or in pairs and occasionally form short chains of up to three cells. Reproduction took place by monopolar budding (apical or sub-apical). (Pseudo)hyphae were not present on GYPA or MEA after up to 12 weeks of incubation. In addition, ascospore formation was not observed in pure cultures or when mixing both strains on GYPA and MEA plates at 25°C for up to 12 weeks. On CHROMagar *Candida* Plus, colonies of both strains appeared pale cream to lavender with a distinctive blue halo surrounding the colony (Figure 3 D3-4). This was the exact same phenotype as the type strain of *C. auris* (CBS 10913^T^, Figure 3 D2). Identification by MALDI-TOF MS did not result in any matched patterns (score values of 0.00–1.69). The closest hits were *C. pseudohaemulonii* (score value 1.56) for CBS 16213^T^ and *Clostridium cadaveris* (score value 1.40) for CBS 16555. To this end, new MALDI-TOF MS main spectra (MSPs) were created for both strains, which can be added to the repository to allow future reliable identification by Bruker Daltonics MALDI-TOF MS. Finally, physiological characteristics were determined and are summarized in Table 1. Compared to close phylogenetic relatives, unique characteristics seem to be the delayed but positive fermentation of maltose, negative growth on 50%–60% glucose, tolerance to 0.1% cycloheximide and growth up to 42°C for both strains.

**Figure 3.**
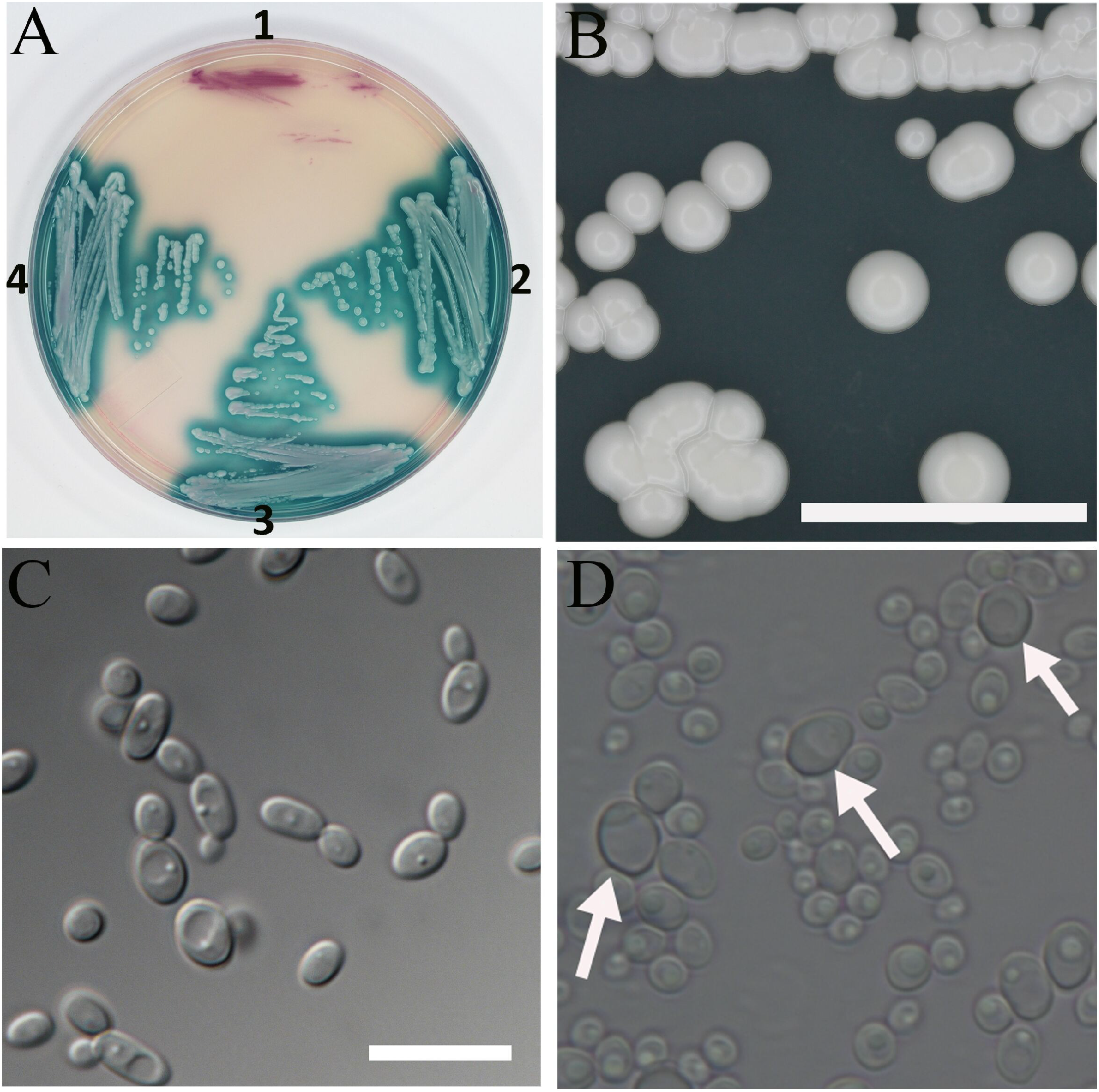
Morphology of strain CBS 16213^T^. (A) colony morphology on GYPA after incubation for 7 days. (B) Yeast cells from an overnight culture in YPD. (C) Yeast cells from a 7-day old culture on GYPA showing a sub-population of larger cells (arrows). (D) Colonies grown on CHROMagar *Candida* Plus: (1) *Candida haemulonii* CBS 5149^T^, (2) *Candida auris* CBS 10913^T^, (3) *Candida khanbhai* CBS 16213^T^, (4) *Candida khanbhai* CBS 16555. Bars, 1 cm for (A), 10 μm for (B, C).

**Table 1.**
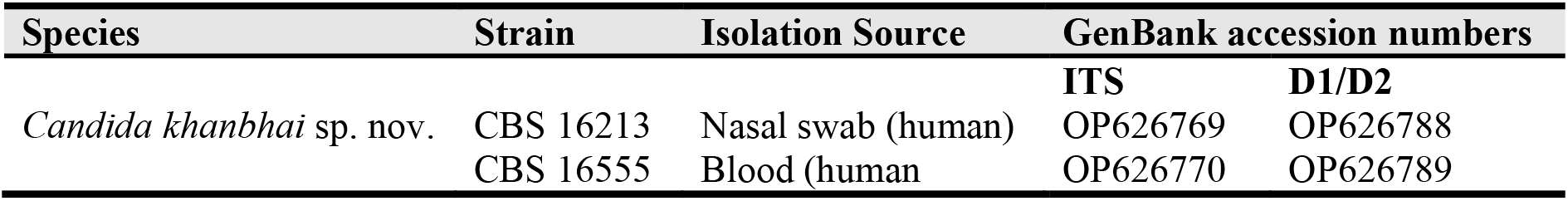
Isolation sources and NCBI GenBank accession numbers of the yeast strains characterized in this study.

| Species | Strain | Isolation Source | GenBank accession numbers |  |
| --- | --- | --- | --- | --- |
|  |  |  | ITS | D1/D2 |
| *Candida khanbhai* sp. nov. | CBS 16213 | Nasal swab (human) | OP626769 | OP626788 |
|  | CBS 16555 | Blood (human) | OP626770 | OP626789 |

### Antifungal susceptibilities

The in vitro antifungal susceptibility of both strains was tested for AMB, 5FC, multiple azoles and two echinocandins. Table 2 shows the MIC results. Both strains had elevated MICs for AMB (2–4 μg/mL). Increased MICs were also observed for most of the azoles, especially after 48h. Notably, CBS 16213^T^ is sensitive to 5FC (0.03–0.125 μg/mL), where CBS 16555 has a strong increased MIC value for 5FC (4–16 μg/mL). Both strains were sensitive to ANI and MCF. However, again moderately elevated MICs for ANI (0.5–1 μg/mL) and MCF (0.25-0.5 μg/mL) were observed for CBS 16555 compared to CBS 16213^T^.

**Table 2.**
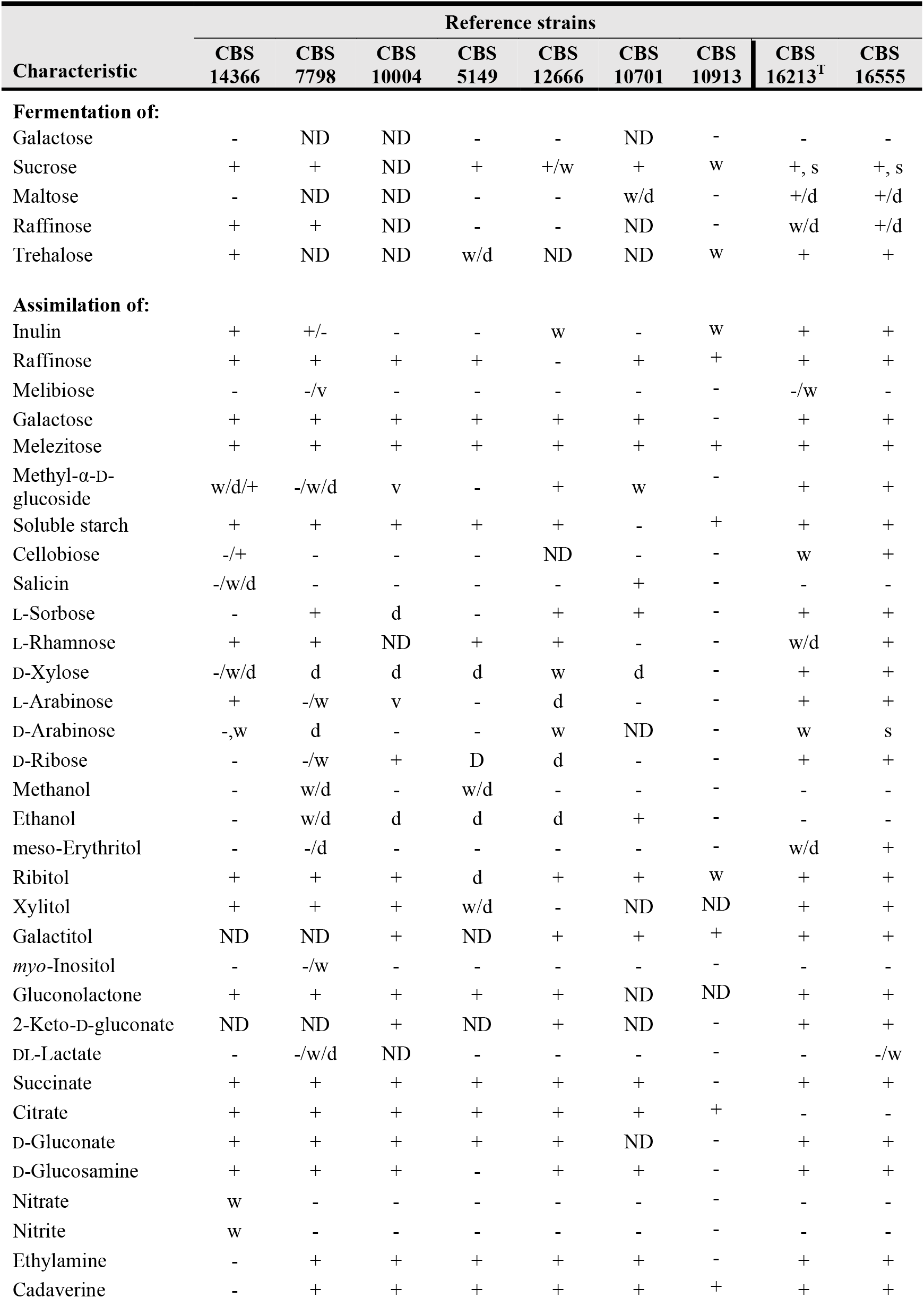
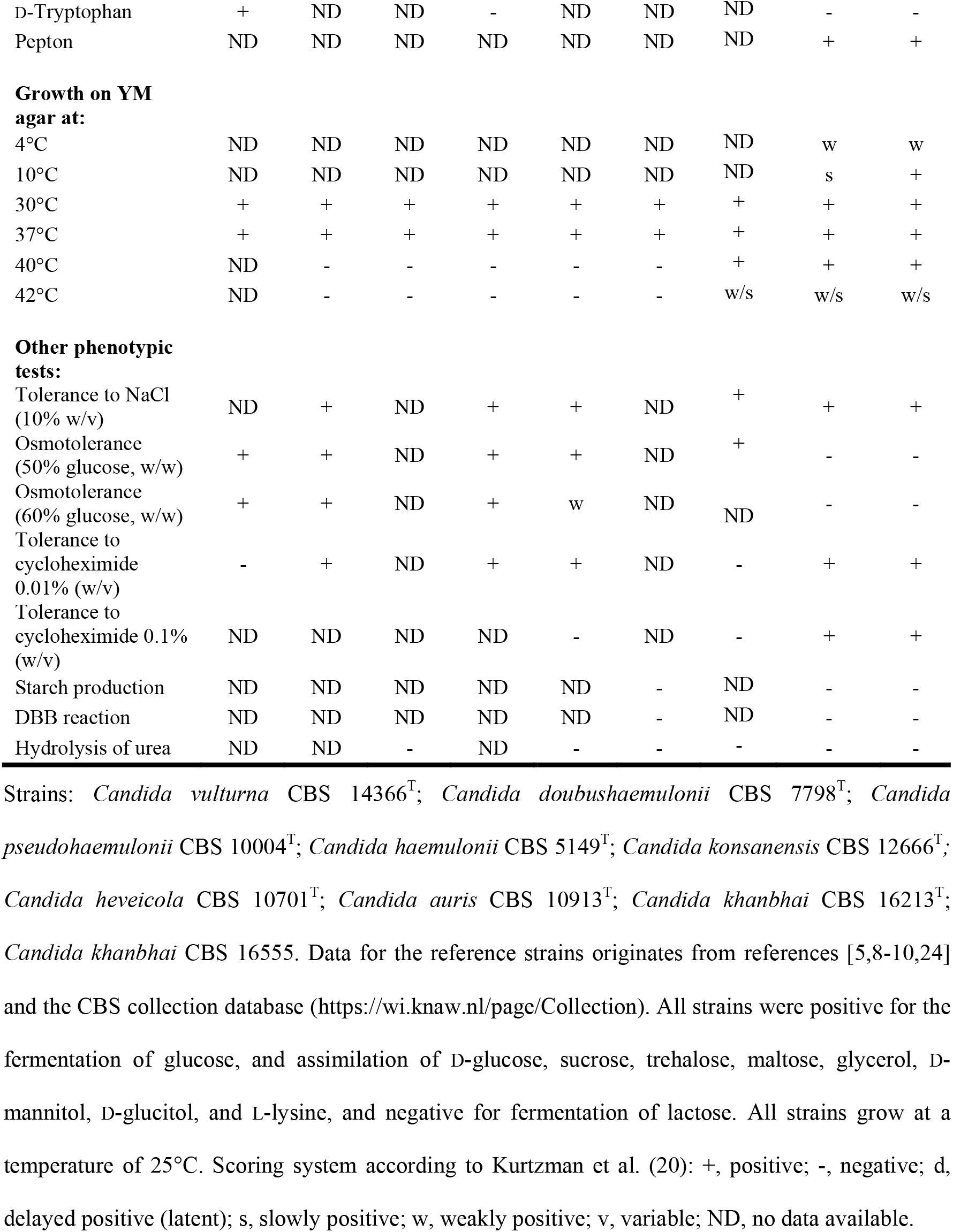
Complete phenotypic characteristics of *Candida khanbhai* strains CBS 16213^T^ and CBS 16555, and their close phylogenetic relatives.

## Discussion

The two new clinical yeast strains in this study came from geographically distinct places. They were isolated from patients in Kuwait (CBS 16213^T^) and Malaysia (CBS 16555). Nevertheless, phylogenetic analysis showed that both strains cluster together and represent a distinct novel species with a close relationship to members of the *C. haemulonii* complex (Figure 1). Although the clinical record of the holotype (CBS 16213) only indicates colonization of the patient, the second strain (CBS 16555) came from a blood culture. The patient infected with strain CBS 16555 passed away despite the use of intravenous fluconazole, suggesting a serious case of resistant candidemia caused by the new species introduced as *C. khanbhai* below. Antifungal susceptibility testing indeed showed reduced sensitivity to multiple antifungal agents. When the tentative MIC breakpoints given for *C. auris* by the CDC are used as references, both strains are resistant to AMB and at least CBS 16555 is resistant to FLU (22). In addition, both strains also showed high MICs for the triazoles tested (Table 2). This is the same multi-resistance pattern as observed in other closely related members of the *C. haemulonii* species complex (5). This resistance together with the ability to infect humans make correct and rapid identification of *C. khanbhai* essential. However, standard biochemical identification systems such as MALDI-TOF MS, VITEK 2 and API 20C AUX misidentified this new species for *C. lusitaniae* and *C. duobushaemulonii*, respectively. Hence, databases should first be updated to enable proper identification of this new pathogenic yeast species. The new CHROMagar *Candida* plus (CCP) is seen as a cost-effective and easy to use alternative to differentiate *C. auris* from its close relatives and more common *Candida* pathogens (23). However, both *C. khanbhai* strains have the exact same phenotype as *C. auris* on this medium (Fig. 1D). Therefore, *C. khanbhai* could cause false positive results on CCP for *C. auris*. On the other hand, possible future infections by *C. khanbhai* are likely to misidentified as *C. auris* when using CCP. Although *C. khanbhai* and *C. auris* do share the same phenotype on CHROMagar and the important trait of growth at 42°C, they have significantly different assimilation patters, which could be used to develop future differentiation assays (Table 1).

Taken together with the description of *C. khanbhai* we add an important clinical *Candida* species to the growing list of emerging fungal pathogens. The isolation from clinical sources at geographically distinct locations together with a worrisome multi-drug resistance pattern and misidentification by commonly used laboratory methods shows many similarities with the story of *C. auris* (6). Therefore, monitoring and correct identification of this species is essential.

### Description of *Candida khanbhai* A.W. de Jong & F. Hagen sp. nov

*Candida khanbhai* (khan.bh ai, ‘khan’ referring to the surname of Prof. dr. Ziauddin Khan who was a medical mycologist, and ‘bhai’ referring to the Hindi word for ‘friend’ and ‘brother’). MycoBank number: MB 846114. Holotype: CBS 16213 preserved in a metabolically inactive state at the CBS culture collection hosted at the Westerdijk Fungal Biodiversity Institute, Utrecht, The Netherlands. Ex-type cultures: CBS 16213, Kw2195/19, 2MG-A0803-17.

On GYPA after 7 days of incubation at 25°C the colonies are off-white coloured, glossy, soft, butyrous and with an entire margin (Fig. 1A). In YPD broth after 1 day of incubation cells are ovoid, ellipsoidal to spherical (1.5–3.5 × 2.5–4.5 μm in diameter), occur singly or in pairs and reproduce by monopolar budding (apical or sub-apical) (Fig 1B, C). On GYPA and MEA (pseudo)hyphae are not formed. Sporulation is not observed in pure cultures or when mixing both strains on GYPA and MEA at 25°C for up to 12 weeks. A summary of the fermentation tests, assimilation tests, and other growth characteristics of strains CBS 16213^T^ and CBS 16555 is provided in Table 1. Minimum inhibitory concentrations of the main classes of antifungal drugs are given in Table 2. The holotype CBS 16213 was isolated in 2019 from a nasal swab of a 73-year-old patient hospitalized in Mubarak Al-Kabir Hospital (Jabriya, Kuwait) and is permanently preserved in a metabolically inactive state in the CBS yeast collection of the Westerdijk Fungal Biodiversity Institute, Utrecht, the Netherlands.

In addition to the description of *C. khanbhai* it was observed that several members of the *Candida haemulonii* species complex were invalidly described. Hence, the opportunity was taken to correct this here, formally validating the names:

*Candida auris* Satoh & Makimura ex F. Hagen sp. nov. MB 846663
For a detailed description see Satoh et al., Microbiol. Immunol. 53 (1): 43 (2009) (reference 9).
Holotype: CBS 10913 preserved in a metabolically inactive state.
Ex-type cultures: JCM15448; CBS10913; DSM21092.
[originally described as: *Candida auris* Satoh & Makimura, Microbiol. Immunol. 53 (1): 43 (2009), nom. inval., Art. 40.7 (Shenzhen)].
*Candida vulturna* Sipiczki & Tap ex F. Hagen sp. nov. MB 846664
For a detailed description see Sipiczki & Tap, Int. J. Syst. Evol. Microbiol. 66(10): 4014 (2016) (reference 8).
Holotype: CBS 14366 preserved in a metabolically inactive state.
Ex-type cultures: CBS 14366; 11-1170; NCAIM-Y02177; CCY 094-001-001.
[originally described as: *Candida vulturna* Sipiczki & Tap, Int. J. Syst. Evol. Microbiol. 66: 4014 (2016), nom. inval., Arts Art. 36.1(b) and 40.7 (Shenzhen)].
*Candida metrosideri* Klaps, C. Vega, C.M. Herrera, Junker, B. Lievens & Álvarez-Pérez ex F. Hagen sp. nov. MB 846665
For a detailed description see Klaps et al., PLoS One 15(10): e0240093, 11 (2020) (reference 25).
Holotype: CBS 16091 preserved in a metabolically inactive state.
Ex-type cultures: CBS 16091; JK22; MUCL 57821.
[originally described as: *Candida metrosideri* Klaps, C. Vega, C.M. Herrera, Junker, B. Lievens & Álvarez-Pérez, PLoS One 15 (10): e0240093, 11 (2020), nom. inval., Art. 36.1(a) (Shenzhen)].
*Candida ohialehuae* Klaps, C. Vega, C.M. Herrera, Junker, B. Lievens & Álvarez-Pérez ex F. Hagen sp. nov. MB 846666
For a detailed description see Klaps et al., PLoS One 15(10): e0240093, 11 (2020) (reference 25).
Holotype: CBS 16092 preserved in a metabolically inactive state.
Ex-type cultures: CBS 16092; JK58.2; MUCL 57822.
[originally described as: *Candida ohialehuae* Klaps, C. Vega, C.M. Herrera, Junker, B. Lievens & Álvarez-Pérez, PLoS One 15 (10): e0240093, 11 (2020), nom. inval., Art. 36.1(a) (Shenzhen)].
*Candida chanthaburiensis* Limtong & Yongman. ex F. Hagen sp. nov. MB 846667
For a detailed description see Limtong & Yongmanitchai, Antonie van Leeuwenhoek 98: 383 (2010) (reference 26).
Holotype: CBS 10926 preserved in a metabolic inactive state.
Ex-type cultures: CBS 10926; EM33; NBRC 102176.
[originally described as: *Candida chanthaburiensis* Limtong & Yongman., Antonie van Leeuwenhoek 98 (3): 383 (2010), nom. inval., Art. 40.7 (Shenzhen)]
*Candida konsanensis* Sarawan, Mahakhan, Jindam., K. Vichitph., S. Vichitph. & Sawaengk. ex F. Hagen sp. nov. MB 846668
For a detailed description, see Sarawan et al., World J Microbiol Biotechnol. 29: 1483 (2013) (reference 24).
Holotype: CBS 10701 preserved in a metabolic inactive state.
Ex-type cultures: CBS 10701; KKU-FW10; BCC 5258; NBRC 109082.
[originally described as: *Candida konsanensis* Sarawan, Mahakhan, Jindam., K. Vichitph., S. Vichitph. & Sawaengk., World J. Microbiol. Biotechnol. 29: 1483 (2013), nom. inval., Arts 40.7, F.5.1 (Shenzhen)]

**Table 3.**
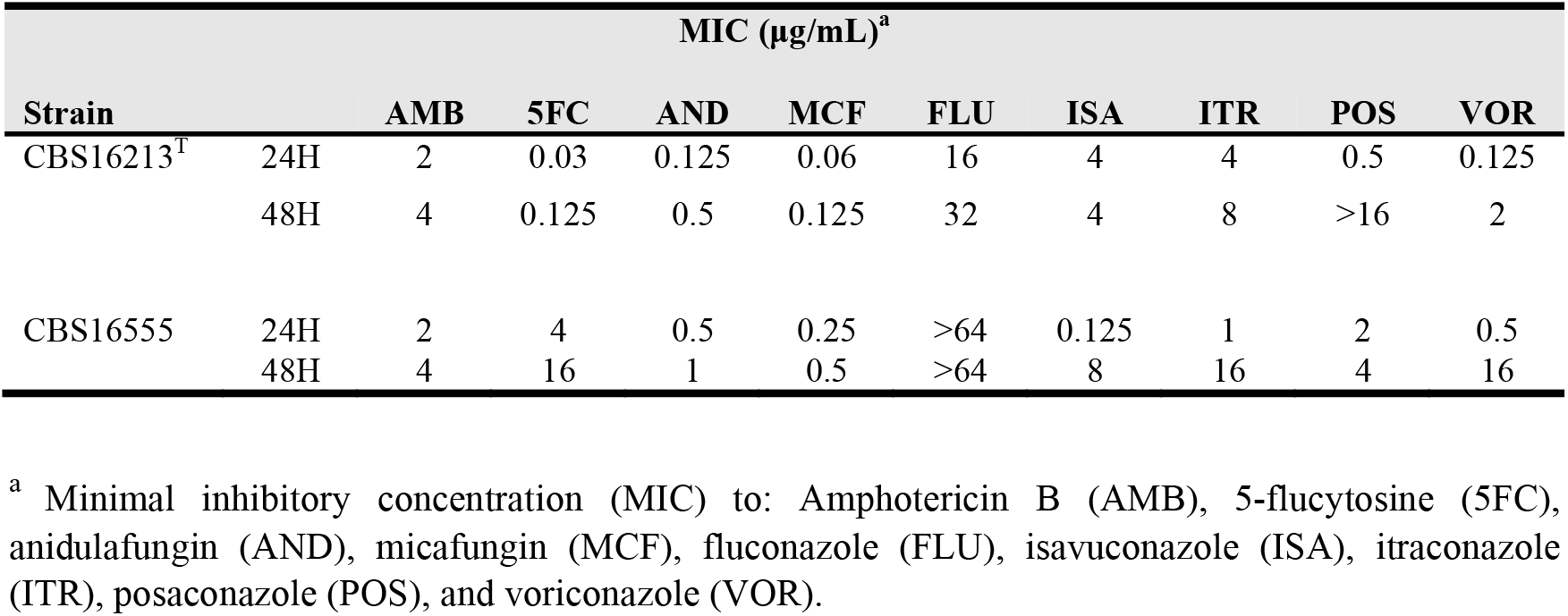
Antifungal susceptibility profiles of strains CBS 16213^T^ and CBS 16555 characterized in this study.

## Acknowledgements

We thank Konstanze Bensch for her valuable input related to the taxonomic status of the here described novel species and formalizing the invalidly described species. The Centers for Disease Control & Prevention (Atlanta, GA, USA) is acknowledged for providing the majority of the *Candida auris* reference strains via the CDC Isolate Bank.

## Data Availability

Yeast strains used in this study have been deposited in the CBS culture collection (hosted at the Westerdijk Fungal Biodiversity Institute, Utrecht, The Netherlands) or are available via the CDC Isolate Bank (hosted at the Centers for Disease Control & Prevention, Atlanta, GA, USA). The generated sequences have been deposited in the NCBI GenBank repository. The strains and sequence data accession numbers are indicated throughout the manuscript, all accession numbers are provided in Figure 2.

